# Psychoneuroimmunology of Viral Hepatitis and Hepatocellular Carcinoma: A Systematic Review and Meta-Analysis

**DOI:** 10.64898/2026.11.04.764298

**Authors:** Adriel Leal Nóbile, Roseane Galdioli Nava, Ronaldy Santana Santos, Fernando Yuri Nery do Vale, Júlia Nakanishi Usuda, Anny Silva Adri, Yohan Lucas Gonçalves Corrêa, Amanda Rodriguez Valera, Luciana Bezerra de Souza Gianasi, Gustavo Cabral-Miranda, Lena F Schimke Marques, Rodrigo J S Dalmolin, Rohini Sharma, Hans Dieter Ochs, Igor Salerno Filgueiras, Haroldo Dutra Dias, Otavio Cabral-Marques

## Abstract

**Objective:** Hepatocellular carcinoma (HCC) and viral hepatitis (VH) are recognized primarily as hepatic diseases, yet they impose a substantial neuropsychiatric disorders (NPDs) encompassing depression, anxiety, cognitive impairment, and fatigue. Whether these manifestations reflect psychological reactions or biologically grounded psychoneuroimmune dysfunction remains poorly understood.

**Methods:** Here, we present a pre-registered systematic review, PRISMA check-list with meta-analysis registration (PROSPERO 2026 CRD420251081757) and risk of bias methods to synthesizing evidence from screened studies combining psychosocial outcomes with immune dysregulation markers across HCC and VH populations.

**Results:** 24 studies were selected for bibliometric and network analyses identified a growing psychoneuroimmunology research hub linking cytokine signaling, microglial activation, blood-brain barrier (BBB) disruption, and gut-liver-brain axis dysregulation to the neuropsychiatric phenotype of these diseases. Meta-analysis revealed a pooled hazard ratio (HR) of 1.92 (95% CI: 1.09–3.39) for adverse outcomes in HCC and a pooled HR of 2.28 (95% CI: 1.48–3.52) for depression associated to genetic variants in VH. TNF-α, but not IL-6, showed a tendency toward lower levels in VH patients with versus without NPDs, suggesting differential inflammatory roles.

**Conclusion:** A psychoneuroimmunological approach, integrating immune, endocrine, neural, and psychosocial dimensions, is essential for capturing the full systemic impact of HCC and VH and should inform future mechanistic research and multidisciplinary clinical strategies, the literature remains fragmented by heterogeneous designs and a paucity of longitudinal data.

## INTRODUCTION

Viral hepatitis (VH) and hepatocellular carcinoma (HCC) represent a connected disease continuum of growing global concern. Hepatitis arises from multiple etiological factors, including poor nutrition, alcohol misuse, substance abuse, autoimmunity and viral infections^1,2^. Among these, hepatitis B virus (HBV), hepatitis C virus (HCV), and hepatitis D virus (HDV) are etiologies strongly associated with chronic hepatic inflammation and progressive liver damage, which can ultimately culminate in HCC, the main and predominant histological subtype of primary liver cancer^2,3^. HCC is the third leading cause of cancer related death worldwide and displays marked geographic heterogeneity^4^. Critically, both VH and HCC extend well beyond the hepatic compartment, manifesting systemically as chronic fatigue, sleep disturbances, hepatic encephalopathy, arthritis, thrombocytopenia, thyroid dysfunction, glomerulonephritis, proteinuria, peripheral neuropathy, and, importantly, an increased risk of cognitive impairment and neuropsychiatric disorders (NPDs)^2,5,6^.

Despite this broad systemic burden, the relationship between HCC, VH and NPDs, including depression, anxiety, cognitive impairment, and schizophrenia, remains insufficiently explored, as in other infections^5^. When neuropsychiatric symptoms are reported in these patients, they are typically interpreted in isolation, with limited consideration of the immunologic mechanisms that may contribute to their development. This reductionist approach overlooks the fundamental principles of psychoneuroimmunology (PNI)^7–9^. These extrinsic factors such as psychological stress, environmental adversity, and social disruption, and intrinsic factors such as immune, nervous, and endocrine systems interactions, are intertwined and may shape both health and behavior of patients affected with HCC or VH^6,7^. This occurs through the interplay of cytokines, systemic inflammation, and NPDs, mechanisms increasingly implicated across a broad spectrum of diseases ^10–12^. Yet the existing literature remains fragmented and predominantly unifactorial, precluding a coherent mechanistic understanding of these associations.

Here, we address this gap through a systematic review and meta-analysis that synthesizes studies combining psychosocial or cognitive outcomes with immune dysregulation markers, focusing on cytokine measurements, in the context of HCC and VH. Our aims are threefold: to characterize the current state and limitations of multidisciplinary PNI related research, particularly with respect to the simultaneous investigation of brain-immune and systemic interactions, the classification of psychiatric disorders, and interdisciplinary diagnostic and therapeutic approaches in these diseases; to quantitatively assess the association between inflammatory biomarkers and neuropsychiatric outcomes; and to evaluate whether such an approach can support more integrated models of clinical care and treatment for patients with VH and HCC.

## METHODS

### Search Strategy

This systematic review was conducted in accordance with PRISMA (Preferred Reporting Items for Systematic Reviews and Meta-Analyses) guidelines and prospectively registered in PROSPERO (CRD420251081757)^13,14^. The primary aim was to synthesize evidence for the association between cellular signaling mechanisms and neuropsychiatric outcomes in HCC and VH.

We based our study on the PECOS strategy: Population (P), individuals with HCC or VH; Exposure (E), cellular signaling mechanisms, including biochemical and cytokine markers; Comparator (C), healthy controls or patients without these conditions or without relevant pathway alterations; Outcomes (O), depression, mood disorders, and related neuropsychiatric symptoms; Study (S) design, as clinical cohort studies, intervention, observational, case-control and cross-sectional^15^. The central question we asked was whether immunological and/or biochemical biomarkers undergo changes that affect neuropsychiatric outcomes during HCC and VH conditions.

Relevant publications were identified through a comprehensive literature search of PubMed/MEDLINE, Embase, Scopus, and EBSCO from database inception to December 2025. The search strategy combined terms related to: “viral hepatitis OR HCC” AND “neuropsychiatric outcomes” AND “psychoneuroimmunological” AND “cellular signaling mechanisms”, including inflammatory and cytokine-related pathways (see Supplementary Materials and Supplementary Table 1-3 for full search terms by database).

### Eligibility Criteria and Reasons for Exclusion

Studies were included if they met the following PECOS criteria: (1) involved patients with HCC or VH, regardless of disease stage or etiology; (2) evaluated psychoneuroimmunological, inflammatory, or cellular signaling mechanisms, including biochemical or cytokine-related markers; and (3) reported neuropsychiatric outcomes, such as depression, anxiety, mood disorders, cognitive impairment, or related psychosocial measures. Observational and clinical studies were considered eligible for meta-analysis. No language restrictions were applied beyond the predefined eligibility criteria.

Neuropsychiatric outcomes were considered when assessed using psychiatric and neurocognitive instruments. For mood and depressive symptoms, we selected the Beck Depression Inventory (BDI and BDI-II)^16^, the Perceived Stress Scale combined with the Patient Health Questionnaire-9 (PSS/PHQ-9)^17^, Major Depressive Episode (MDE) and the Mini International Neuropsychiatric Interview (MINI) based on Diagnostic and Statistical Manual of Mental Disorders, 5th ed. (DSM-5)^18^. Cognitive outcomes were considered when studies used the Stanford-Binet or more specific measures such as the Ruff Figural Fluency Test (RFFT)^19^, the California Verbal Learning Test (CVLT)^20^, and the Grooved Pegboard^21^, which assess executive function, verbal memory, and psychomotor performance, respectively. Overall, the included studies combined psychiatric diagnostic tools, self-report scales, and neuropsychological tests to characterize NPDs outcomes in HCC and VH.

The following publications were excluded: studies not involving HCC or VH; studies not addressing psychoneuroimmune and/or cellular signaling mechanisms; articles without clearly defined mental health or neuropsychiatric outcomes; *in vitro*-only studies; conference abstracts; editorials; letters; narrative reviews; and previously published systematic reviews or meta-analyses. Studies were also excluded when full texts were unavailable or when the data were insufficient for extraction. Bioinformatics, case reports and animals models were not included in the main synthesis, but were considered to support the interpretation and discussion of the findings.

### Study Screening and Data Extraction

Two reviewers (A.L.N and F.Y.N.V) independently screened the titles and abstracts of all retrieved records to identify potentially eligible studies. Full-text articles were then assessed independently according to predefined inclusion and exclusion criteria. Duplicate removal and the screening process were conducted blindly using the Rayyan platform to minimize selection bias^22^. Disagreements were resolved through discussion and, when necessary, by consultation with a third reviewer (I.S.F). Inter-reviewer agreement during study selection was quantified using Cohen’s Kappa coefficient^23^.

Data were extracted independently by the same reviewers using a standardised spreadsheet. After duplication removal, for each study, we collected information on publication year, country, sample size, study design, population characteristics, VH or HCC condition, NPDs outcomes, psychometric or cognitive assessment tools, immune or cellular signaling markers, and the main findings related to psychoneuroimmunological mechanisms for bibliometric analysis (**See Supplementary Materials**). Where applicable, information on control groups, clinical stage, cytokine and marker measures, and treatment status was also recorded. When data were unclear or incomplete, they were checked against the full text and supplementary materials. The entire review process was quality assessed using the PRISMA checklist (**Supplementary Table 2**).

For neuropsychiatric outcomes, we extracted both categorical and continuous measures, including formal psychiatric diagnoses, depressive symptom scores, stress-related measures, and cognitive test performance. Biomarker data included cytokines, inflammatory mediators and other cellular signaling indicators relevant to HCC, VH, and neuropsychiatric comorbidities, like serum albumin, alanine aminotransferase (ALT), alkaline phosphatase (ALP) and aspartate aminotransferase (AST). From the study tables, we extracted values reported as interquartile range (IQR), standardized mean difference (SMD), viral load, and demographic variables such as age and sex, as well as markers of liver integrity, prioritising ALT, AST, lactate dehydrogenase (LDH), and alpha-fetoprotein.

### Quality Assessment and Risk of Bias

The Risk of bias was independently assessed by two reviewers (A.L.N and R.G.N) according to the study design and the corresponding gold-standard appraisal tool. For longitudinal/prospective and prospective translational studies that compared clinical interventions or treatment regimens in real-world settings, the ROBINS-I (Cochrane) tool was applied^24^. For analytical cross-sectional and case-control studies, we used the Joanna Briggs Institute (JBI) Checklist for Analytical Cross-Sectional Studies and JBI Checklist for Case-Control Studies^25^, with particular attention to the adequacy of control selection and comparability of exposure measurement between cases and controls (**See Supplementary Materials and Supplementary Table 5-7**)^35^ **Statistical Analysis** Statistical analyses were performed when the data reported in the selected articles were sufficient to allow quantitative synthesis. Serum biomarker data were analyzed using SMD meta-analysis and correlation-based synthesis, depending on the structure of the available data^26,27^. For cytokine concentrations, values were first harmonized to common units and, when necessary, converted from median and interquartile range into approximate mean and variance.

An ordinal severity variable was then defined to represent three groups: G1) VH with NPDs, G2) VH without NPDs, and G3) healthy controls. For each cytokine, an ordinal moderator and study-level clustering were included as a random effect. This approach allowed the to estimation of both baseline cytokine levels in healthy controls and the change in concentration across increasing levels of clinical treatments. Single-arm forest plots were generated to visualize study-level estimates and pooled random-effects summaries, together with heterogeneity statistics including I² and τ² ^28^.

To examine small-study effects and dispersion of cytokine results, funnel plots were constructed both at the cytokine level and at the aggregated study level^29^. For pairwise cytokine comparisons, SMD were calculated after reconstructing variance estimates from reported medians and interquartile ranges, and these effect sizes were used to visualise asymmetry as a function of standard error. In addition, correlation coefficients between cytokines and NPDs outcomes were transformed using Fisher’s z test and synthesised with random-effects models^30^. These correlation-based results were then back-transformed to Pearson’s r and further explored using heatmaps summarising the direction, magnitude, and statistical significance of associations across cytokines, psychiatric domains, viruses, and studies.

Given heterogeneity across study designs, populations, and reported outcomes, extracted data were synthesized qualitatively and, when feasible, quantitatively via meta-analysis. Quantitative synthesis was performed when at least three studies evaluated the same biomarker, allowing direct comparison across studies; otherwise, single-arm analyses were conducted. This approach was applied to biochemical markers associated with liver function and systemic inflammation, including ALT, AST, and serum albumin, using SMDs to estimate effect sizes across groups^28^.

All statistical analyses are detailed on supplementary materials and were conducted in R (version 4.5.2) using RStudio^31^, and the analytical scripts are publicly available at the following GitHub repository: https://github.com/adrielleal/article_psychoneuroimmunology_hcc_and_vh.

### Artificial intelligence tool

In this article, the artificial intelligence tool Rayyan was used to remove duplicate articles. The authors declare that they have fully reviewed the content and assume full responsibility for all information presented. This statement is in accordance with the journal recommendations.

## RESULTS

### Systematic identification of psychoneuroimmunological studies in HCC and VH

A comprehensive search across four databases identified 3,714 records, comprising 1,600 studies related to HCC and 2,114 studies focused on VH and associated NPDs. Following duplicate removal, 323 for HCC and 529 VH studies, and blinded title/abstract screening based on PECOS criteria, 29 HCC studies and 39 VH studies were considered eligible for full-text assessment.

However, studies that could not be accessed in full text due to regional and country specific access policies were excluded, as well as for institutional and industrial access requirements, or ethical data protection constraints, as well as those for which eligibility could not be resolved by consensus after third reviewer adjudication, resulting in the exclusion of 22 HCC and 15 VH reports prior to eligibility confirmation. Inter-reviewer agreement score was high for the HCC screening process (Cohen’s κ = 0.94), eliminating the need for third-reviewer arbitration and yielding three studies for inclusion. By contrast, agreement was substantially lower for VH studies (κ = 0.27), reflecting greater heterogeneity in study design and outcome reporting. After third-reviewer adjudication, 20 VH plus one hand-searched article were included in the final synthesis, totalizing 21 studies (**Supplementary Table 3**). The complete selection workflow is summarized in the PRISMA diagram (**See Supplementary Methods and Supplementary Figure 1 for full details**).

### Bibliometric analyses reveal the rapid emergence of psychoneuroimmunology in liver disease research

Temporal bibliometric analyses demonstrated a pronounced expansion of studies investigating NPDs in the context of both HCC and VH, particularly after the early 2000s (**Figure 1a**). Although HCC-related publications consistently predominated throughout the analyzed period, VH-associated studies exhibited accelerated growth after 2020, progressively reducing the disparity between both research fields. This temporal convergence parallels the broader expansion of PNI research during the 21st century, reflecting increasing recognition that emotional, neurological, and immune dysregulations are deeply interconnected across chronic infectious and neoplastic diseases^5,32–36^.

**Figure 1.**
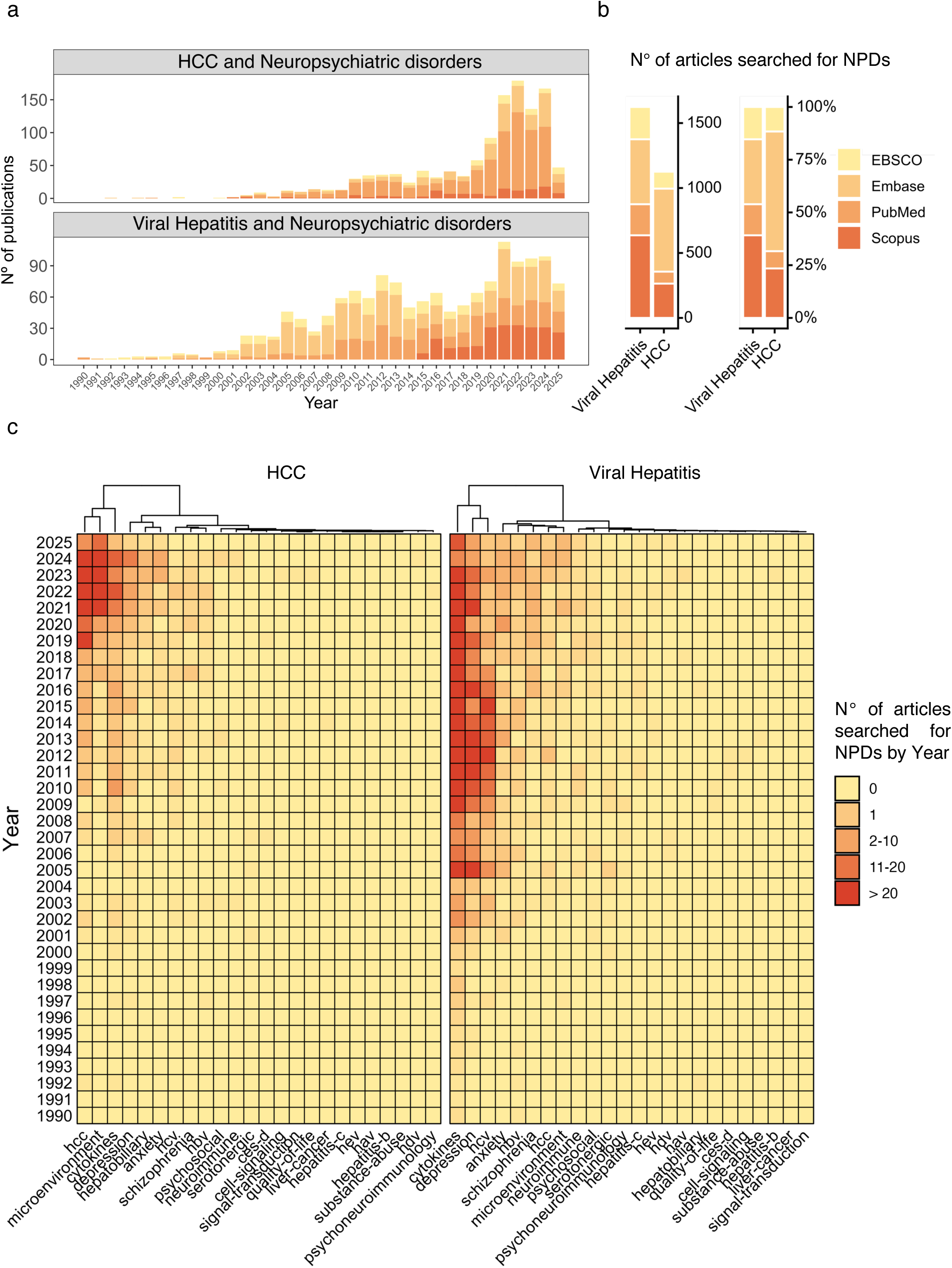
Bibliometric analysis of the search and screened articles. a) Bibliometric analysis indicates a marked temporal growth in publications addressing HCC and VH in association with NPDs, with a more pronounced expansion in the VH literature. Stacked bar plots show that publication output increased substantially after the turn of the 21st century, in PubMed, Embase, Scopus, and EBSCO b) with viral hepatitis-related studies consistently outnumbering those focused on HCC. c) Heatmaps further demonstrate that research activities became more concentrated over time in domains such as depression, anxiety, fatigue, mental health, cognition, and cytokines, especially from the 2010s onward. HCC literature appears more sparse and clustered around later years, whereas in VH the distribution is broader and more sustained across multiple thematic domains.

Database contribution analyses further revealed that Embase and Scopus accounted for the majority of retrieved studies, whereas PubMed and EBSCO contributed proportionally fewer publications to the final data (**Figure 1b**). This distribution suggests that investigations linking neuropsychiatric manifestations to liver disease remain dispersed across multidisciplinary and non-traditional biomedical indexing platforms, reflecting the intrinsically cross-disciplinary nature of PNI research.

After duplicate removal in Rayyan, keyword temporal mapping revealed a marked emergence and expansion of terms associated with psychoneuroimmune dysfunction after the year 2000, including *depression*, *cytokines*, *anxiety*, *schizophrenia*, *psychoneuroimmunology*, *fatigue*, and *microenvironment* (**Figure 1c** and **Supplementary Table 3**). Notably, *depression* and *cytokines* emerged as dominant and progressively convergent concepts across both HCC and VH datasets, suggesting that inflammatory-neuropsychiatric interactions have become central organizing themes in contemporary liver disease research.

While HCC studies demonstrated stronger enrichment for tumor-associated terminology, VH investigations exhibited broader incorporation of neuropsychiatric and immune-inflammatory descriptors over time. These findings support the emergence of a systems-level conceptual plan in which chronic liver diseases are increasingly investigated through integrated psychoneuroimmune and behavioral dimensions rather than as isolated hepatic disorders alone.

### Global bibliometric mapping identifies convergent psychoneuroimmune signatures across HCC and VH research

Comparative term-frequency analyses revealed *depression* and *cytokines* as the most dominant and recurrent concepts across both HCC and VH neuropsychiatric literature (**Figure 2a**), reinforcing the central role of inflammatory-neuropsychiatric interactions in chronic liver disease research. Notably, cytokine associated terminology represented the single most enriched category in both conditions, appearing in 32.8% of VH studies and 21.2% of HCC investigations, whereas *depression* emerged as the predominant neuropsychiatric descriptor, occurring in 19.8% and 11.9% of VH and HCC studies, respectively.

**Figure 2.**
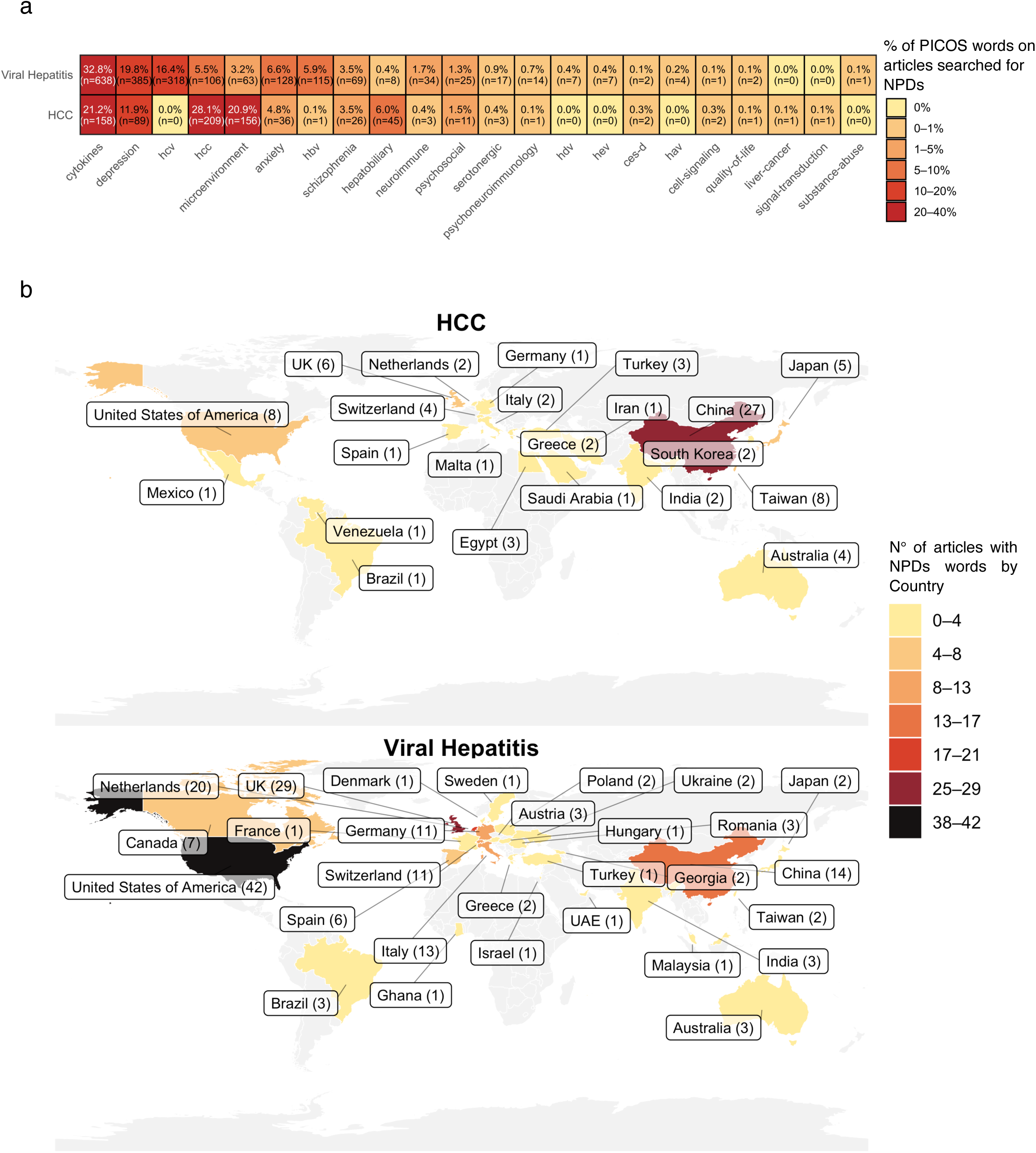
Bibliometric analysis of PECOS used words and NPDs associated studies geographic distribution. a) Heatmap shows differences in PECOS words distribution of the literature on HCC and VH in relation to NPDs. In both fields, the most frequently represented topics were cytokines, and depression, although these domains were proportionally more prominent in VH than in HCC. HCC-related studies showed greater relative contribution of terms such as HCC and PNI, whereas VH studies were more strongly concentrated on classic psychiatric and behavioral domains, including depression and anxiety. b) World maps further demonstrate a distinct geographic pattern: research on HCC was predominantly concentrated in China, with additional contributions from Taiwan, the United States, the United Kingdom, and Japan, while the VH literature was more geographically dispersed, with the United States, United Kingdom, Netherlands, China, Italy, and Germany representing major contributors.

The analysis of the ’cross-contamination’ keywords revealed that the overlap between both search domains remained minimal, with HBV related terms appearing in only 0.1% of HCC-screened studies, supporting the specificity and discriminatory robustness of the PECOS-based curation strategy. HCC-related studies were strongly enriched for tumor-centered terms, particularly *HCC* itself (28.1%) and *microenvironment* (20.9%), consistent with the growing incorporation of immune-oncology into hepatocarcinogenesis research. By contrast, VH investigations displayed stronger virus infections descriptors, especially *HCV* (16.4%) and *HBV* (5.9%), reflecting the infectious and epidemiological orientation of the research in chronic hepatitis, when compared to more acute infections related to VH, *HEV* accounts for 0.4%, and *HAV* for 0.2%.

Geospatial analyses further demonstrated that investigations linking NPDs to liver disease constitute a globally distributed research effort (**Figure 2b**). HCC-related neuropsychiatric studies were predominantly concentrated in East Asia, particularly China, reflecting the high regional burden of HCC. Contrariwise, VH neuropsychiatric research exhibited broader international distribution, with major contributions from North America and Europe, especially the United States, United Kingdom, and the Netherlands. Collectively, these findings reveal that PNI has evolved into a transnational and interdisciplinary research spanning oncology, virology, immunology, inflammatory-neuropsychiatric, and systems medicine.

### Network analyses identify geographically structured psychoneuroimmunology hubs in liver disease research

Integrated keyword co-occurrence and geospatial analyses revealed marked regional organization within the PNI literature associated with liver disease (**Figure 3a**). Studies linking NPDs to HCC were predominantly concentrated in East Asia and North America, with China and the United States emerging as major research epicenters. Yet VH neuropsychiatric investigations displayed a substantially broader international distribution encompassing North America, Europe, the Middle East, Latin America, and East Asia, suggesting that VH associated PNI research has evolved within a more globally interconnected scientific ecosystem. These regional asymmetries may reflect differences in epidemiological burden, healthcare infrastructure, translational oncology priorities, and national research funding strategies^2,4^.

**Figure 3.**
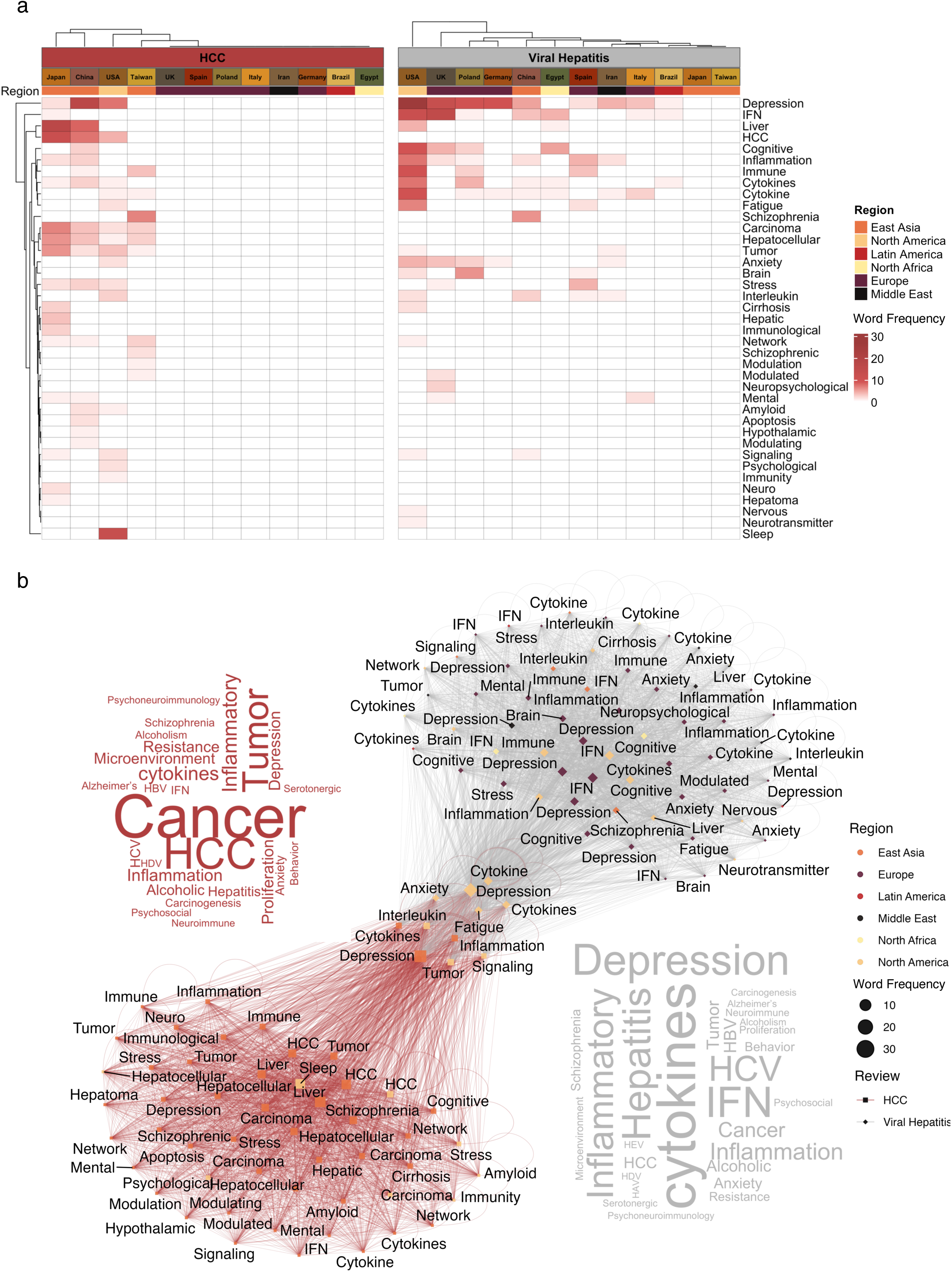
PECOS search keywords characterization using titles and abstracts. a) Clear differences in the regional and thematic structure of the literature on HCC and VH related to NPDs. In the heatmap, both topics share recurrent themes such as depression, IFN, cytokines, inflammation, cognition, fatigue, and liver-related terms, but their distribution across regions is distinct. In HCC, the literature is more concentrated in East Asia, particularly China, Japan, and Taiwan, with additional but smaller contributions from the United States and a few European countries. The VH literature is more geographically dispersed, with stronger contributions from North America, Europe, and East Asia, especially the United States, United Kingdom, Germany, Italy, Spain, and China. b) Word clouds highlight the words that are hubs in a network, “cytokines, depression, interleukin, fatigue, and inflammation”, that reinforce differences of HCC and VH field, dominated by disease-centered and tumor-related terms such as HCC, cancer, tumor, hepatocellular, liver, inflammation, cytokines, and depression, whereas the viral hepatitis field is centered more strongly on depression, cytokines, HCV, hepatitis, IFN, inflammation, and broader neuropsychiatric constructs including anxiety, cognition, fatigue, and stress.

Network and Wordcloud topology analyses further uncovered a highly interconnected PNI interaction hub bridging immune-inflammatory and neuropsychiatric domains (**Figure 3b**). Central nodes included *cytokines*, *inflammation*, *interleukin signaling*, *immune* and *tumor* terms, which densely co-occurred with neurobehavioral descriptors such as *depression*, *anxiety*, *fatigue*, *stress*, *cognitive*, and *neurotransmitter*. Notably, studies from North America and East Asia demonstrated the network centrality for psychoneuroimmune signatures, potentially indicating scientific investment by major global research economies in deciphering the mechanistic interplay between chronic inflammation, neural dysfunction, and psychiatric manifestations in VH and HCC.

### Heterogeneous study designs constrain adjusted evidence linking liver disease to neuropsychiatric outcomes

Across the 24 included studies, three focused on HCC and 21 on VH. Although these studies collectively examined the relationship between liver disease and NPDs, most were not designed to generate directly comparable adjusted effect estimates. Adjusted odds ratios (OR), risk ratio (RR), or hazard ratio (HR) estimates were rarely reported (**Supplementary Table 4**)^32,37–39^. Only four studies adjusted for common comorbidities, such as hypertension and diabetes, including two HCC and two VH focused studies. HCC studies mainly reported HR for survival-related outcomes, whereas VH studies reported OR for interferon-induced depression. After hand-searching, only one extra study was identified for VH group, Jen Wang L, 2017, which did not report OR, RR, or HR estimates but had cytokines measuares^40^.

This lack of harmonization extended to neuropsychiatric assessment. Three of the 24 studies did not use formal psychometric evaluation, and the others relied on different diagnostic or cognitive tools, including Center for Epidemiologic Studies Depression Scale (CES-D), Hospital Anxiety and Depression Scale (HADS), Diagnostic and Statistical Manual of Mental Disorders (DSM) based diagnostic codes, and cognitive testing batteries.

Excluding the hand-searched article, bioinformatics studies, and case reports, which were used only for discussion, the risk of bias assessment was applied to 23 studies, previolsy had-searching: ROBINS-I (9 studies), JBI case report (2 studies), and JBI case-control (12 studies)(**See Supplementary Materials, Supplementary Figure 1-3, Supplementary Tables 8-11**, **for the full curation and risk of bias analyses results**).

After risk of bias assessment, only a subset of the available evidence could be taken forward into quantitative synthesis . Of the 24 included studies, only seven (30%) provided effect estimates suitable for meta-analysis, two of three HCC studies and five of 21 VH studies, and a few had OR and HR metrics (**Supplementary Tables 12**). This limited pool reflected the broader structure of the field: most studies were biologically informative, but few were designed to generate harmonized risk estimates across neuropsychiatric, inflammatory, biochemical, and viral outcomes^34,41–44^.

Only the VH search group was eligible for comparative cytokine evaluation, which included six studies with comparative measures (**Supplementary Tables 13 and 14**), four studies assessing biochemical markers (**Supplementary Table 15**), and three studies viral markers (**Supplementary Table 16**)^45,52–55^

Despite this restriction, the pooled analyses revealed clinically relevant associations in both disease contexts. In HCC, the random-effects model showed a pooled HR of 1.92 (95% CI: 1.09–3.39), indicating that psychological and clinical factors, including the diagnosis of depression, CES-D scores, and vascular invasion, were associated with poorer outcomes. In VH, the pooled OR was 2.28 (95% CI: 1.48– 3.52), suggesting that genetic variants previously implicated in interferon-α-induced depression and serotonergic signalling were associated with higher odds of depression-related outcomes **(Supplementary Figure 3).** These effects may indicate that neuropsychiatric vulnerability is not merely a secondary clinical annotation in liver disease, but may intersect with prognosis, treatment response, and host inflammatory susceptibility^39,51^

### Cytokine dysregulation emerges as the dominant inflammatory signal linking viral hepatitis to neuropsychiatric outcomes

Two-arm cytokine meta-analyses suggested that individual cytokines behaved differently reinforced by the heterogeneity analysis (**Figure 4a and 4b, Supplementary Figure 4, Supplementary Table 13**). IL-6 showed a pooled SMD close to zero (SMD = − MDed SMDary Table 13 4,lyses suggested that individual cytokines behaved differently reinforced by the heterogeneity analysis (t may intersect with prognosis, treatment response, and host inflammatory ng that psychological and clinicinflammatory status in VH rather than a marker that reliably distinguishes neuropsychiatric status (**Figure 4c**). TNF-α showed a stronger and more directional pattern, with a pooled negative estimate (SMD = −0.51, 95% CI: −1.10 to 0.07), although the confidence interval crossed zero (**Figure 4d**). This tendency was driven by heterogeneous study-level effects: Bahramabadi et al.^43^ reported a moderate negative effect in HBV-positive patients, Cunha et al.^42^ showed a significant negative association in HCV-positive patients, and Hilsabeck et al.^36^ showed no meaningful difference.^45,55^

**Figure 4.**
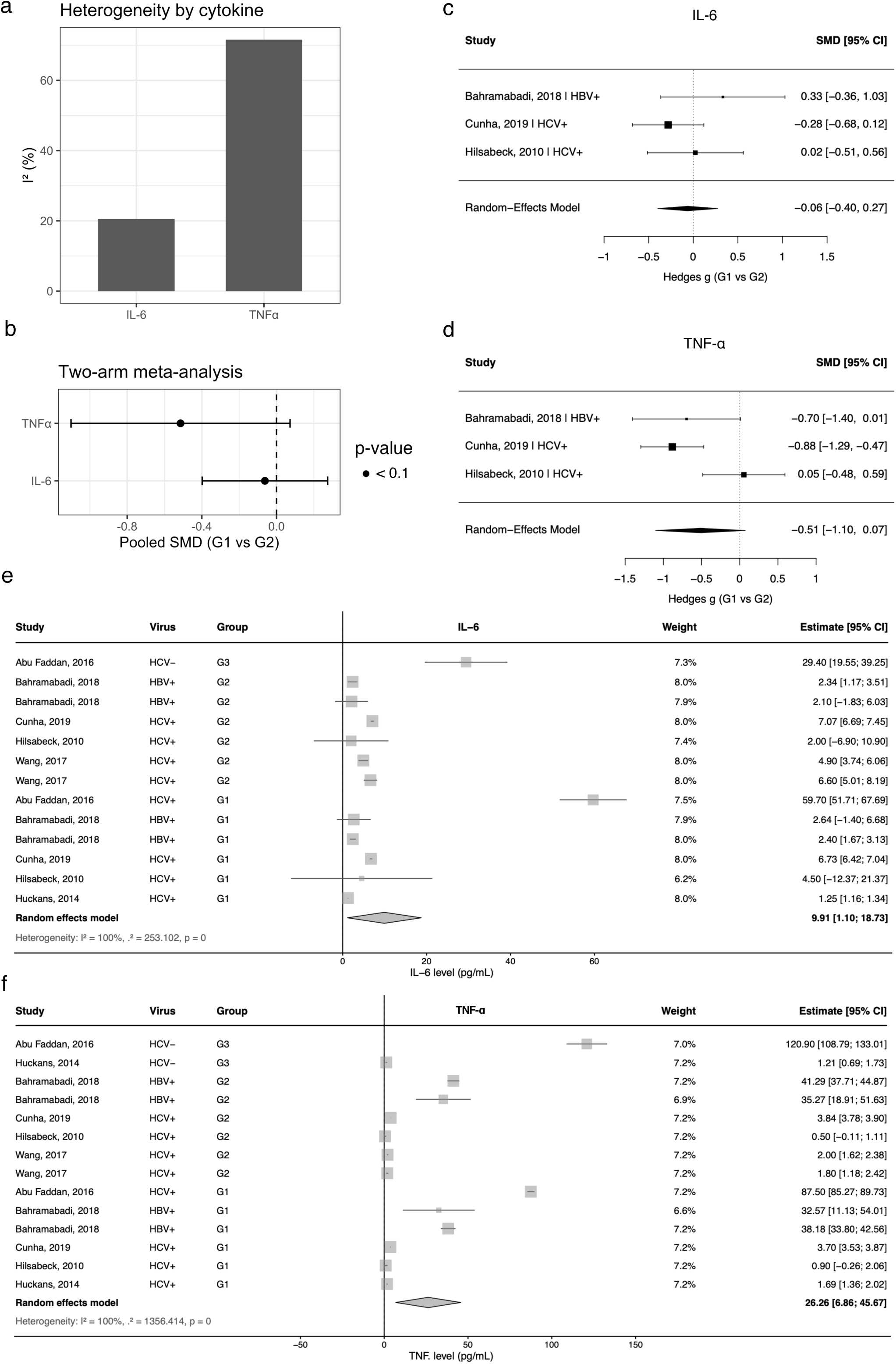
**Heterogeneity assessment and meta-analytical synthesis of IL-6 and TNF-**α **in VH studies**. a) Bar plot showing the degree of between-study heterogeneity for the two cytokines included in the quantitative synthesis, expressed as I² (%), indicating lower heterogeneity for IL-6 and substantially higher heterogeneity for TNF-α. b) Two-arm meta-analysis comparing G1 (viral hepatitis with NPDs) versus G2 (viral hepatitis without NPDs) using standardized mean differences (SMDs), summarizing the pooled^36,42,43^ effects for IL-6 and TNF-α and their 95% confidence intervals. c) Forest plot of the single-arm meta-analysis for IL-6, presenting individual study estimates, weights, and the pooled random-effects estimate across G1, G2, and healthy controls^36,42,43^. d) Forest plot of the single-arm meta-analysis for TNF-α, showing individual study estimates, weights, and the pooled random-effects estimate across the same groups^36,42,43^. e) Funnel-plot–based evaluation of IL-6 as a candidate for quantitative synthesis, illustrating the distribution of effect sizes and standard errors across studies^36,40,42,43,45,53^. f) Funnel-plot–based evaluation of TNF-α, used to visually assess between-study dispersion, asymmetry, and the potential suitability of this cytokine for meta-analysis^36,40,42,43,45,53^.

In addition, single-arm meta-analyses indicated that both IL-6 and TNF-α were elevated across VH cohorts. However, these estimates were not consistent across studies, cytokine levels varied substantially between cohorts and clinical groups, as reflected by very high heterogeneity (I² = 100% for both cytokines). The pooled estimate for IL-6 was 9.91 pg/mL (95% CI: 1.10–18.73), whereas the pooled estimate for TNF-α was 26.26 pg/mL (95% CI: 6.86–45.67). Some of the highest cytokine levels occurred in HCV-related cohorts, particularly in the report by Abu Faddan et al.^45^, which contributed substantially to the dispersion of the pooled estimates.TNF-α appeared more sensitive than IL-6 to NPD variation, but its effect was context dependent rather than uniform across VH cohorts (**Figure 4e and 4f**).

Correlation pattern of HCV associated to NPDs among studies provides the strongest support for the observation that cytokine associations were not confined to depression alone, nor did they onto a single neuropsychiatric domain. Instead, inflammatory markers were distributed across depression, anxiety, cognition, personality-related traits, fatigue, mental health, physical health, and work productivity^34–36,44–46^. TNF-α and IL-8 showed recurrent associations with depression, IL-8 appeared prominently in anxiety-related outcomes, and TNF-α, IL-6, IL-8, and IFN-γ were linked to cognition and personality-related domains (**Figure 5a and Supplementary Table 14**).

**Figure 5.**
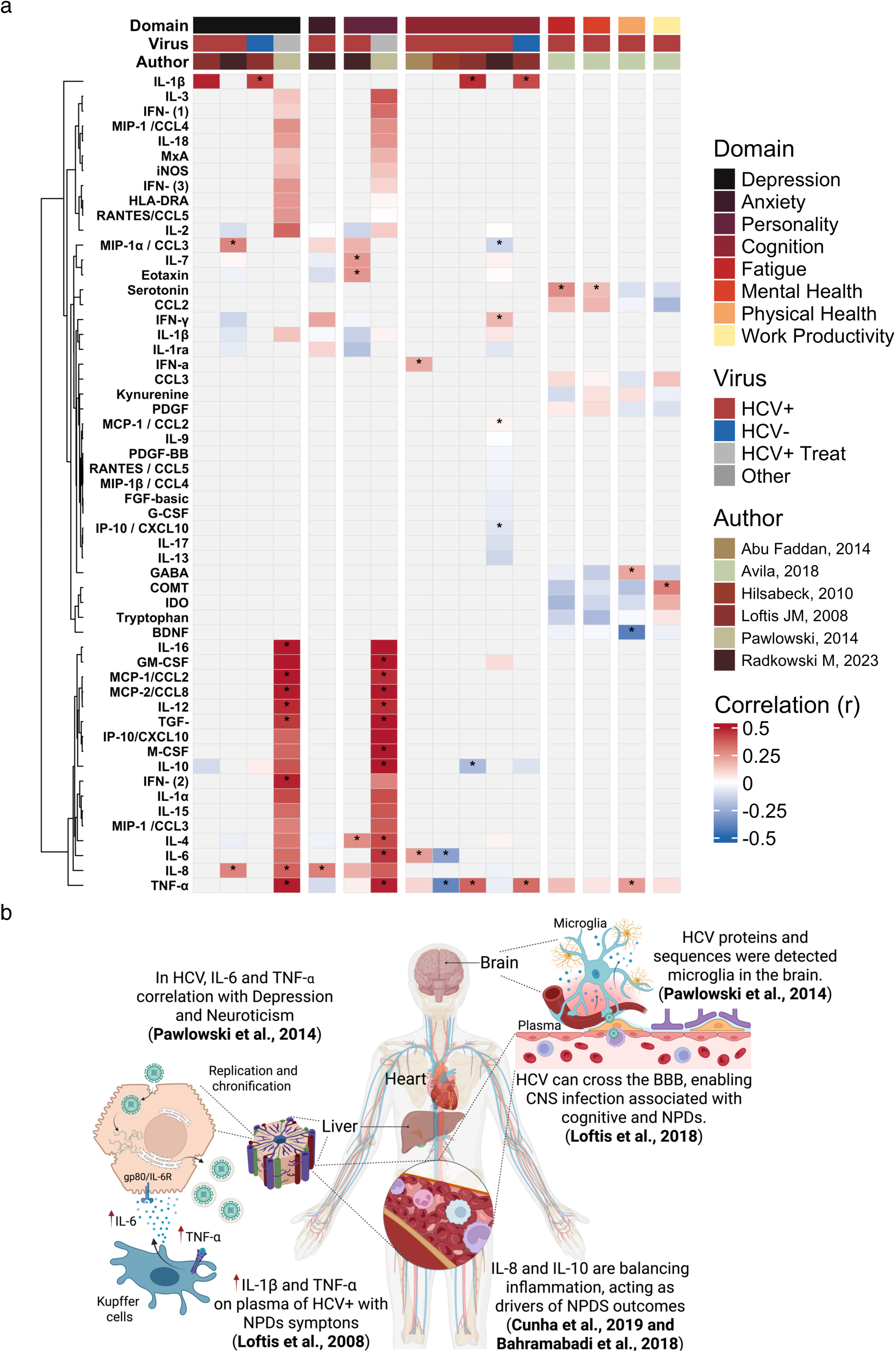
Cytokines correlations and main functions on NPDs and neuroimmune burden. **(a)** Correlation heatmap based on Fisher’s *z* transformation, showing the direction and magnitude of associations between cytokines and NPD domains across the included viral hepatitis studies. The heatmap relates cytokines, TNF-α, IL-8, IL-6, and IFN-γ, to NPDs domains such as depression, anxiety, personality, cognition, fatigue, mental health, physical health, and work productivity, stratified by viral status and study author. Positive correlations are shown in red tones and negative correlations in blue tones^34–36,44–46^. **(b)** Mechanistic overview of the selected viral hepatitis studies, summarizing the main hypothesized pathways linking HCV infection to NPDs, including hepatic replication and chronic inflammation, Kupffer cell activation, systemic cytokine release, blood-brain barrier crossing, microglial involvement, and central nervous system effects^34,41–44^.

By contrast, conventional liver and viral markers were less informative for determining NPD status. HCV viral load varied widely across studies, and the pooled random-effects estimate showed substantial heterogeneity, indicating that viral burden did not consistently distinguish patients with NPDs from those without^36^. Similarly, serum albumin remained relatively stable, whereas ALT and AST were frequently elevated but highly variable^42,45^(**Supplementary Figure 5**). Thus, although viral replication and liver injury remain important features of disease activity, they do not appear to independently explain the neuropsychiatric phenotype.

Integrating these findings, HCV replication and chronic hepatic inflammation may activate Kupffer cells and peripheral immune cells, amplifying systemic cytokine release^44^. These inflammatory signals may interact with the central nervous system (CNS) through blood-brain barrier (BBB) disruption, viral protein or sequence exposure, microglial activation, and peripheral immune-cell communication^34,41^. Thus, the neuropsychiatric burden observed in HCV infection may reflect not a single peripheral marker, but a distributed inflammatory axis connecting hepatic injury, circulating immunity, and brain function (**Figure 5b**). Within this context, VH may shape neuropsychiatric burden through persistent inflammatory communication between the liver, peripheral immune cells, endothelial barriers, and the CNS.

### Psychoneuroimmunological Mechanisms as Therapeutic Opportunities in Hepatocellular Carcinoma

Of the seven HCC studies include (**Supplementary Table 8**), only Jozuka et al. explicitly applied a PNI intervention, although limited to a single case, providing an important longitudinal clinical example in which psychiatric symptoms, biochemical markers and immune mediators were combined for HCC management. The patient had HCC with major depressive disorder and received a combined psychoneuroimmunological treatment strategy, surviving for more than five years after diagnosis^47^.

Steel et al demonstrated associations between cytokine activity, sleep disturbance, and clinical outcomes in patients with HCC^32^. Consistent with these observations, Zhao et al. showed in a murine Chronic Unpredictable Mild Stress (CUMS) model that chronic stress-related exposures, including social isolation and sleep disruption, were associated with increased PD-1 expression and enhanced tumor progression (**Figure 6a**)^48^. Although direct human-mouse comparisons are not possible (**Figure 6b**), this study suggests that depressive symptoms were accompanied by the normalization of biochemical markers of liver disease severity from pathological to healthy levels over time (**Figure 6c**), as well as in cytokine levels, including IL-2, IL-6, IL-12, and IFN-γ (**Figure 6d and 6e**).

**Figure 6.**
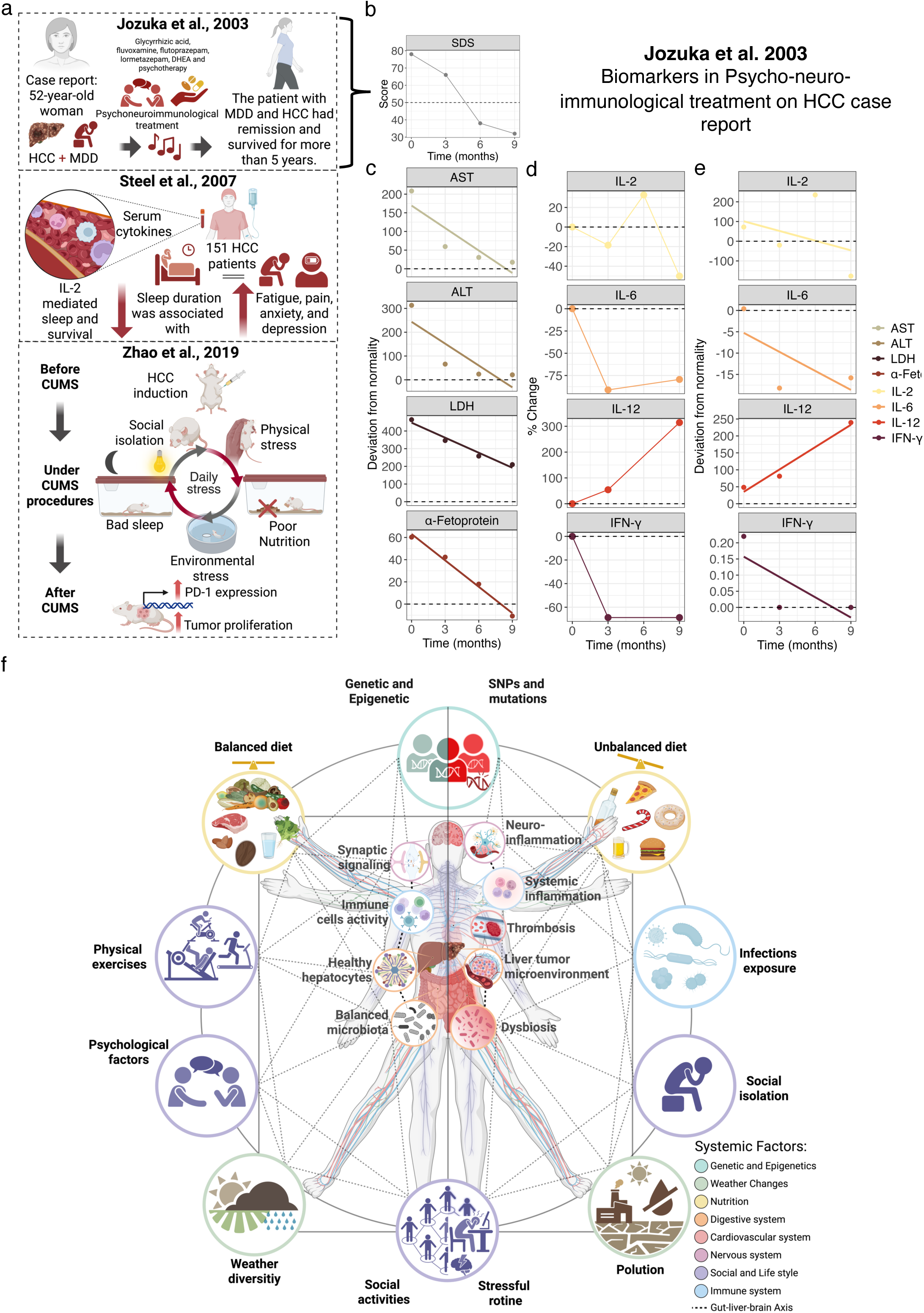
HCC relations to NPDs symtons. (a-e) Exploratory summary of the findings from Jozuka et al.^32,47,48^ showing the longitudinal evolution of depressive symptoms, SDS score, liver biochemical markers, AST, ALT, LDH, α-fetoprotein, and cytokines, IL-2, IL-6, IL-12, and IFN-γ, over follow-up, including both percentage change and deviation from normal values. (f) Integrative overview of the reviewed literature on VH and HCC, summarizing the main systemic, behavioral, environmental, immunological, and neuroinflammatory pathways potentially linking liver disease to neuropsychiatric outcomes. The scheme highlights factors such as diet, infections, social isolation, stress, pollution, genetic and epigenetic influences, microbiota imbalance, liver tumor microenvironment, thrombosis, systemic inflammation, neuroinflammation, synaptic signaling, and immune-cell activity within a proposed gut-liver-brain axis structure^32–36,40–47,53–55,60^.

Collectively, psychoneuroimmune interactions in HCC and VH may influenced by a broad range of biological, behavioral, and environmental factors. Diet, sleep quality, physical activity, psychosocial stress, social isolation, and environmental exposures may modulate gut-liver-brain communication and systemic immune responses, highlighting the multidimensional nature of psychoneuroimmunological research processes (**Figure** 6f)^10,11,40^.

## DISCUSSION

This systematic review and meta-analysis supports cytokine dysregulation as the most consistent inflammatory signal linking VH and HCC to NPDs. Thus, although depression and related symptoms are commonly associated with a pro-inflammatory state, in HCC and VH they are unlikely to represent merely emotional responses to malignancy, systemic inflammation, or chronic infection. Rather, they may reflect a biologically embedded psychoneuroimmune burden arising from the interaction between tumour biology, hepatic dysfunction, immune response, and psychosocial stress^10,49^.

Our findings support the need to investigate VH and HCC through psychoneuroimmunological approachs^5,7,9^, rather than as an exclusively hepatic condition by integrating the immune, endocrine, neural, psychosocial, and nutritional dimensions that may jointly shape disease expression, treatment response, and quality of life^6^. Across the included studies, neuropsychiatric manifestations such as depression, anxiety, cognitive impairment, fatigue, schizophrenia, and stress-related symptoms emerged alongside alterations in inflammatory, immune, and neuroendocrine signaling, extending beyond liver inflammation in both VH^34–36,45,50^ and HCC^33,47,48,51,52^.

A recurring theme throughout the literature has been the role of immune dysregulation as a plausible mechanistic link between hepatic pathophysiology and neuropsychiatric symptoms^52,53^. Cytokines such as IL-6, TNF-α, IL-2, and IFN-γ were repeatedly implicated, together with endocrine mediators and markers of systemic stress^10,35–37,40,41,43,45,46,54,55^. In VH, particularly HBV and HCV, chronic immune activation may contribute to fatigue, depressive symptoms, and cognitive dysfunction even before overt tumor development^35,36,43^.

In HCC, these mechanisms may be further amplified by tumor induced inflammation, hepatic dysfunction, and the cumulative psychosocial burden of advanced disease, including sleep disturbance, all of which may adversely affect survival. In parallel, inflammatory mediators such as IL-6, IL-8, and TNF-α may help link HCC to NPD outcomes, either independently of or in interaction with IFN immune pathways^32,37^.

The available evidence does not support a single-cytokine or single-marker model. Instead, the data point to a multidimensional immune signature in which TNF-α, IL-6, IL-8, IL-1β, IL-10, and IFN-γ participate in a liver-immune-brain axis^40^. Within this context, VH may promote neuropsychiatric vulnerability through sustained inflammatory communication between hepatic tissue, peripheral blood immune cells, endothelial barriers, and the CNS^41^. Although substantial heterogeneity and limited availability of adjusted estimates require cautious interpretation, the convergence between mechanistic evidence, pooled cytokine analyses, and correlation patterns indicates that inflammatory network disruption may be more closely aligned with NPD outcomes than with viral load or conventional biochemical markers.

The reviewed evidence further suggests that chronic VH, particularly HCV infection similar to interferon treated diseases, may create a sustained inflammatory environment that predisposes patients to later NPD vulnerability, while in HCC additional systemic disruption occurs through tumor progression, altered liver metabolism, reduced physiological resilience, and glucocorticoid-mediated PD-1 upregulation^33,38,40^.

Specific biological phenomena, such as the ability of HCV to cross the BBB^41^, may lead to infection of the CNS and the generation of inflammatory cytokines. This hypothesis is supported by the observation that HCV proteins can be detected in brain microglia^34^ and may contribute to neurobehavioral deficits, including mild cognitive impairment, fatigue, depression, and psychosis, in these diseases^34,41^. The resulting neuropsychiatric symptoms are further modified by overlapping mechanisms including inflammatory signaling, another unexplored layers of neuroendocrine dysregulation, and by advanced liver disease, hepatic encephalopathy^6,49^. Together, these observations reinforce the importance of considering VH and HCC as part of a continuum in which immune-mediated alterations influence both psychiatric outcomes and overall disease severity.

From a translational perspective, these findings highlight the need for a multidisciplinary approach to patient care. Routine psychiatric and cognitive screening appears not to be fully integrated into hepatology and oncology guidelines, despite evidence that these neuropsychiatric symptoms affect quality of life, adherence to treatment, and possibly long term prognosis^5,40^. A psychoneuroimmunological perspective may therefore improve patient stratification by combining clinical outcomes with biomarkers of inflammation and cellular signaling, and by monitoring patients for infection, cancer and NPDs. Although the current literature remains heterogeneous, it provides a rationale for future studies aimed at identifying psychoneuroimmune signatures that could support early recognition of vulnerable patients and guide integrated interventions.

Jozuka et al. applied PNI intervention, combining psychotherapy, antidepressant and anti-inflammatory therapy, music, and aromatherapy in an HCC patient with major depression, providing an illustrative example, the patient becoming symptom free, of this integrative approach^47^. However, this uncontrolled single case report should be interpreted cautiously without claiming direct relationship with cancer remission^24^. Nevertheless, the longitudinal monitoring of depressive symptoms, biochemical markers, and cytokines highlights the type of multidimensional assessment that future controlled studies should develop more rigorously^24,47^.

Bioinformatic studies also offer an important exploratory layer for this type of studies. Although they cannot replace clinical validation, they may help identify transcriptional dysregulation, protein targets, and candidate “neuroimmunome” pathways relevant to VH and HCC. The studies included in the discussion point to dysregulated networks involving immune signaling, glutamate receptors, neurotransmission, and pathways associated with depression, Alzheimer’s disease, schizophrenia, disease severity, and survival. However, these findings should be interpreted as hypothesis-generating rather than definitive, although they provide a useful basis for prioritizing these aspects in future translational studies^33,51,52,56^.

The included studies varied substantially in design, sample size, psychiatric assessment methods, biomarker selection, disease stage, viral status, treatment exposure, and confounder adjustment, putting limitations on comparisons. In addition, adjusted OR, RR, or HR estimates were uncommon, and only a subset of studies could be included in quantitative synthesis. These limitations underscore the need for longitudinal, adequately powered, and mechanistically informed studies that integrate psychiatric phenotyping with immune, endocrine, neurological, microbial, biochemical, and clinical trajectories, an integration that surpasses the classic gut-liver-brain axis investigation, and their clinical trajectories across both VH and HCC populations^49,57–59^.

Overall, this systematic review suggests that the neuropsychiatric burden observed in VH and HCC is closely linked to PNI mechanisms. Rather than representing secondary or isolated complications, these manifestations may be intrinsic to the systemic biology of chronic liver disease and liver cancer. Advancing this field will require closer integration among psychiatry, hepatology, endocrinology, oncology, immunology, neurology, and systems medicine. Such integration would be challenging, but would enable the translation of psychoneuroimmune mechanisms into clinically meaningful tools for diagnosis, monitoring, risk stratification, and integrative treatment.

### Ethic statement

Not applicable.

### Author Contributions

A.L.N.: Data curation, Methodology, Formal analysis, Visualization, Writing – original draft. R.G.N. and R.S.S: Conceptualization, Methodology, Writing – review & editing. F.Y.N.V., J.N.U., A.S.A., Y.L.G.C., A.R.V., L.B.S.G., G.C.M., L.F.S.M., R.J.S.D., R.S., H.D.O., I.S.F., and H.D.D.: Writing – review & editing. O.C.M.: Conceptualization, Supervision, Writing – review & editing.

## Acknowledgments

We thank the São Paulo Research Foundation (FAPESP grants 2018/18886-9 to O.C.M, 2023/12268-0 to A.S.A, 2023/14417-2 to J.N.U, 2024/08016-8 to A.L.N, 2025/07090-2 to F.Y.N.V and 2024/22162-7 to R.G.N) for financial support. We acknowledge the National Council for Scientific and Technological Development (CNPq) (grants: 309482/2022-4 to O.C.M and 130027/2023-5 to Y.L.G.C). We acknowledge the grant CNPq/SECTICS/CAPES/FAPs N° 46/2024, process 408662/2024-7 to O.C.M for financial support. We acknowledge the “Coordenação de Aperfeiçoamento de Pessoal de Nível Superior - Brasil (CAPES)” for financioal support 88887.082794/2014-00 to R.S.S and 88887.196113/2025-00 to I.S.F. The Article Processing Charge for the publication of this research was funded by CAPES”.

## Data Availability Statement

The data that support the findings were downloaded from EBSCO, Embase, PUBMED and Scopus. Also, the .ris data used in Rayyan, the R scripts used in all analysis are openly available in GitHub repository https://github.com/adrielleal/article_psychoneuroimmunology_hcc_and_vh.

## References

1. Shah, N. D. & Barritt, A. S. Nutrition as Therapy in Liver Disease. Clin. Ther. 44, 682–696 (2022).

2. Odenwald, M. A. & Paul, S. Viral hepatitis: Past, present, and future. World J. Gastroenterol. 28, 1405 (2022).

3. Stella, L., Santopaolo, F., Gasbarrini, A., Pompili, M. & Ponziani, F. R. Viral hepatitis and hepatocellular carcinoma: From molecular pathways to the role of clinical surveillance and antiviral treatment. World J. Gastroenterol. 28, 2251 (2022).

4. Singh, S. P., Madke, T. & Chand, P. Global Epidemiology of Hepatocellular Carcinoma. J. Clin. Exp. Hepatol. 15, 102446 (2025).

5. Yolken, R. Infections and neuropsychiatric disorders: new studies document pathways to prevention and treatment. Mol. Psychiatry 28, 2624–2626 (2023).

6. Cacoub, P. & Asselah, T. Hepatitis B Virus Infection and Extra-Hepatic Manifestations: A Systemic Disease. American Journal of Gastroenterology 117, 253–263 (2022).

7. Yazdanpanah, N. & Rezaei, N. Introduction to Psychoneuroimmunology. Integrated Science 30, 1–16 (2024).

8. Ader, R. Historical Perspectives on Psychoneuroimmunology. *Psychoneuroimmunology*, Stress, and Infection 1–24 (2020) doi:10.1201/9780367812522-1.

9. John B. Jemmott, I. Psychoneuroimmunology. American Behavioral Scientist 28, 497–509 (1985).

10. Nava, R. G. et al. Modulation of neuroimmune cytokine networks by antidepressants: implications in mood regulation. Translational Psychiatry *2025* 15:*1* 15, 314-(2025).

11. Wessa, C., Simon, M. S. & De Picker, L. Current evidence on immune-driven depression. Curr. Opin. Psychiatry 39, 8–18 (2026).

12. Adri, A. S. et al. Systems-level transcriptomic analysis reveals synapse-related gene dysregulation in peripheral leukocytes of MDD patients. Scientific Reports *2026* 16:*1* 16, 8336-(2026).

13. Schiavo, J. H. PROSPERO: An International Register of Systematic Review Protocols. Med. Ref. Serv. Q. 38, 171–180 (2019).

14. Parums, D. V. Editorial: Review Articles, Systematic Reviews, Meta-Analysis, and the Updated Preferred Reporting Items for Systematic Reviews and Meta-Analyses (PRISMA) 2020 Guidelines. Med. Sci. Monit. 27, e934475–1 (2021).

15. Amir-Behghadami, M. & Janati, A. Population, Intervention, Comparison, Outcomes and Study (PICOS) design as a framework to formulate eligibility criteria in systematic reviews. Emergency Medicine Journal 37, 387 (2020).

16. Beck, A. T., Steer, R. A. & Brown, G. Beck Depression Inventory–II. PsycTESTS Dataset https://doi.org/10.1037/T00742-000 (2011) doi:10.1037/T00742-000.

17. Cohen, S. & Janicki-Deverts, D. Who’s Stressed? Distributions of Psychological Stress in the United States in Probability Samples from 1983, 2006, and 2009. J. Appl. Soc. Psychol. 42, 1320–1334 (2012).

18. DIAGNOSTIC AND STATISTICAL MANUAL OF DSM-5 ^TM^.

19. Ruff, R. Figural Fluency Test. Encyclopedia of Clinical Neuropsychology 1–3 (2017) doi:10.1007/978-3-319-56782-2_2048-2.

20. Delis, D. C., Kramer, J. H., Kaplan, E. & Ober, B. A. California Verbal Learning Test--Second Edition. PsycTESTS Dataset https://doi.org/10.1037/T15072-000 (2016) doi:10.1037/T15072-000.

21. Bryden, P. J. & Roy, E. A. A new method of administering the Grooved Pegboard Test: Performance as a function of handedness and sex. Brain Cogn. 58, 258– 268 (2005).

22. Johnson, N. & Phillips, M. Rayyan for systematic reviews. J. Electron. Resour. Librariansh. 30, 46–48 (2018).

23. Vieira, S. M., Kaymak, U. & Sousa, J. M. C. Cohen’s kappa coefficient as a performance measure for feature selection. 2010 IEEE World Congress on Computational Intelligence, WCCI https://doi.org/10.1109/FUZZY.2010.5584447 (2010) doi:10.1109/FUZZY.2010.5584447.

24. Hinneburg, I. ROBINS-1: A tool for asssessing risk of bias in non-randomised studies of interventions. Med. Monatsschr. Pharm. 40, 175–177 (2017).

25. Santos, W. M. Dos, Secoli, S. R. & Püschel, V. A. de A. The Joanna Briggs Institute approach for systematic reviews. Rev. Lat. Am. Enfermagem 26, e3074 (2018).

26. Welz, T., Doebler, P. & Pauly, M. Fisher transformation based confidence intervals of correlations in fixed- and random-effects meta-analysis. British Journal of Mathematical and Statistical Psychology 75, 1–22 (2022).

27. Andrade, C. Mean Difference, Standardized Mean Difference (SMD), and Their Use in Meta-Analysis: As Simple as It Gets. J. Clin. Psychiatry 81, 20f13681–20f13681 (2020).

28. Singh, J., Abrams, K. R. & Bujkiewicz, S. Incorporating single-arm studies in meta-analysis of randomised controlled trials: a simulation study. BMC Medical Research Methodology *2021* 21:*1* 21, 114-(2021).

29. Sterne, J. A. C. & Harbord, R. M. Funnel Plots in Meta-analysis. Stata J. 4, 127– 141 (2004).

30. Welz, T., Doebler, P. & Pauly, M. Fisher transformation based confidence intervals of correlations in fixed- and random-effects meta-analysis. British Journal of Mathematical and Statistical Psychology 75, 1–22 (2022).

31. Posit Software, P. Posit team (2025). RStudio: Integrated Development Environment for R. Preprint at http://www.posit.co/.

32. Steel, J. L., Geller, D. A., Gamblin, T. C., Olek, M. C. & Carr, B. I. Depression, immunity, and survival in patients with hepatobiliary carcinoma. Journal of Clinical Oncology 25, 2397–2405 (2007).

33. Han, T., Zhou, Y. & Li, D. Relationship between hepatocellular carcinoma and depression via online database analysis. Bioengineered 12, 1689–1697 (2021).

34. Pawlowski, T. et al. Depression and neuroticism in patients with chronic hepatitis C: Correlation with peripheral blood mononuclear cells activation. Journal of Clinical Virology 60, 105–111 (2014).

35. Radkowski, M., et al. Depression and Cognitive Dysfunction in Patients with Chronic Hepatitis C: Correlation with Viral Replication in the Peripheral Blood Mononuclear Cells and Cytokines in Serum. Int. J. Mol. Sci. 24, (2023).

36. Hilsabeck, R. C. et al. Cognitive efficiency is associated with endogenous cytokine levels in patients with chronic hepatitis C. J. Neuroimmunol. 221, 53–61 (2010).

37. Steel, J. L. et al. Prospective analyses of cytokine mediation of sleep and survival in the context of advanced cancer. Psychosom. Med. 80, 483–491 (2018).

38. Kraus, M. R. et al. Serotonin-1A Receptor Gene HTR1A Variation Predicts Interferon-Induced Depression in Chronic Hepatitis C. Gastroenterology 132, 1279–1286 (2007).

39. Smith, A. K. et al. Association of a polymorphism in the indoleamine-2,3-dioxygenase gene and interferon-α-induced depression in patients with chronic hepatitis C. Mol. Psychiatry 17, 781–789 (2012).

40. Jen Wang, L., et al. Mental health, pro-inflammatory cytokines, and brain derived neurotrophic factor in patients undergoing treatment for hepatitis C: comparison between peginterferon-alpha-2a and peginterferon-alpha-2b. Neuropsychiatry 07, (2017).

41. Loftis, J. M. et al. S100B and Inflammatory Cytokine Levels in Blood as Potential Markers of Blood–Brain Barrier Damage and Psychiatric Impairment in Comorbid Hepatitis C Viral Infection and Alcohol Use Disorder. Alcohol. Clin. Exp. Res. 42, 1466–1475 (2018).

42. Cunha, L. R. da et al. Interleukin-10 promoter gene polymorphisms are associated with the first major depressive episode in chronic hepatitis C patients. Clin. Res. Hepatol. Gastroenterol. 43, 417–426 (2019).

43. Bahramabadi, R. et al. Serum levels of IL-6, IL-8,TNF-α, and TGF-β in Chronic HBV-infected patients: Effect of depression and anxiety. Lab Medicine 49, 41–46 (2018).

44. Loftis, J. M., Huckans, M., Ruimy, S., Hinrichs, D. J. & Hauser, P. Depressive symptoms in patients with chronic hepatitis C are correlated with elevated plasma levels of interleukin-1β and tumor necrosis factor-α. Neurosci. Lett. 430, 264–268 (2008).

45. Abu Faddan, N. H., et al. Cognitive function and endogenous cytokine levels in children with chronic hepatitis C. J. Viral Hepat. 22, 665–670 (2015).

46. de Avila, L. et al. Cytokine balance is restored as patient-reported outcomes improve in patients recovering from chronic hepatitis C. Liver International 39, 1631–1640 (2019).

47. Jozuka, H., Jozuka, E., Suzuki, M., Takeuchi, S. & Takatsu, Y. Psycho-neuro-immunological treatment of hepatocellular carcinoma with major depression - A single case report. Curr. Med. Res. Opin. 19, 59–63 (2003).

48. Zhao, Y. et al. Depression Promotes Hepatocellular Carcinoma Progression through a Glucocorticoids Mediated Up-Regulation of PD-1 Expression in Tumor infiltrating NK Cells. https://doi.org/10.1093/carcin/bgz017/5306310 doi:10.1093/carcin/bgz017/5306310.

49. Zou, J., Li, J., Wang, X., Tang, D. & Chen, R. Neuroimmune modulation in liver pathophysiology. Journal of Neuroinflammation *2024* 21:*1* 21, 188-(2024).

50. Xue, Y., Sun, X., Li, Y., Liu, X. & Dong, C. Increased risk of hepatitis E virus infection in schizophrenia. Arch. Virol. 158, 359–365 (2013).

51. Wang, Y., Yang, Y., Liang, C. & Zhang, H. Exploring the Roles of Key Mediators IKBKE and HSPA1A in Alzheimer’s Disease and Hepatocellular Carcinoma through Bioinformatics Analysis. Int. J. Mol. Sci. 25, (2024).

52. Huang, K. C. et al. Analysis of schizophrenia and hepatocellular carcinoma genetic network with corresponding modularity and pathways: Novel insights to the immune system. BMC Genomics 14, (2013).

53. Huckans, M. et al. Multi-analyte profile analysis of plasma immune proteins: Altered expression of peripheral immune factors is associated with neuropsychiatric symptom severity in adults with and without chronic hepatitis C virus infection. Brain Behav. 4, 123–142 (2014).

54. Cheng, H. H. et al. Socioeconomic disparities of depressive symptoms and cytokines in hepatocellular carcinoma. Psychooncology 28, 1624–1632 (2019).

55. Falasca, K. et al. Quality of life, depression, and cytokine patterns in patients with chronic hepatitis C treated with antiviral therapy. Clinical and Investigative Medicine 32, (2009).

56. Nóbile, A. L. et al. The Neuroimmunome of Hepatitis Patients Associates With Disease Severity. J. Med. Virol. 97, (2025).

57. Kronsten, V. T., Tranah, T. H., Pariante, C. & Shawcross, D. L. Gut-derived systemic inflammation as a driver of depression in chronic liver disease. Journal of Hepatology vol. 76 665–680 Preprint at 10.1016/j.jhep.2021.11.008 (2022).

58. Lieb, K. et al. Cognitive impairment in patients with chronic hepatitis treated with interferon alpha (IFNα): results from a prospective study. European Psychiatry 21, 204–210 (2006).

59. Ponce-Regalado, M. D. et al. NeuroImmunoEndocrinology: A brief historic narrative. J. Leukoc. Biol. 112, 97–114 (2022).

60. Wang, Y., DelRocco, N. & Lin, L. Comparisons of various estimates of the I2 statistic for quantifying between-study heterogeneity in meta-analysis. Stat. Methods Med. Res. 33, 745–764 (2024).

